# Influence of stimulus complexity on the specificity of visual perceptual learning

**DOI:** 10.1101/832303

**Authors:** Shahab Bakhtiari, Asmara Awada, Christopher C. Pack

## Abstract

Although the structure and function of the human visual system are determined in large part during early development, there is ample evidence for adult plasticity as well. Such plasticity has important consequences for restoring vision after cortical damage and for improving function in healthy people. Although these applications have shown promising results, they are often limited by pathological *specificity*: Improvements obtained through perceptual training fail to generalize beyond the trained stimulus feature or location. Efforts to reduce specificity have focused on the design of training tasks, but less is known about the effects of stimulus structure on the specificity of perceptual learning. Here, we leverage physiological findings from the dorsal visual pathway of the primate brain to explore the hypothesis that learning specificity is related to the complexity of the training stimulus. Specifically, because neurons in higher-level structures of the dorsal visual pathway exhibit little stimulus specificity, we reasoned that training with more complex stimuli would reduce the specificity of learning. We trained human observers on stimuli of varying complexity, ranging from simple sinewave gratings to complex optic flow fields. Our results show that training with more complex stimuli reduces specificity for spatial position and stimulus features. Such changes are associated with increased spatial integration. These findings were captured by a computational “reweighting” model that decoded the outputs of simulated neurons in areas MT and MST of the primate visual cortex. Our results suggest that the addition of more complex stimuli into perceptual learning paradigms provides a simple and effective way to minimize specificity in learning.

## Introduction

The visual system is highly plastic during early life (Hubel, Wiesel, LeVay, Barlow, & Gaze, 1977), and recent work has demonstrated substantial plasticity in adults as well. Specifically, the field of visual perceptual learning has shown that training leads to changes in the ability to discriminate orientations, colors, shapes, and other stimulus features (Goldstone, 1998; Seitz & Watanabe, 2005). However, to be of practical utility, the effects of perceptual learning should generalize beyond the stimuli used during training, and as a result there is considerable research devoted to the question of the *specificity* (or conversely, the *transfer*) of perceptual learning (Ahissar & Hochstein, 1997; Jeter, Dosher, Petrov, & Lu, 2009).

In all likelihood, the specificity of perceptual learning is related to the hierarchical organization of the visual cortex: Low-level cortical areas, such as V1, exhibit exquisite specificity for low-level stimulus features, such as orientation and spatial position. Higher-level structures, in contrast, are often selective for stimulus features such as shape or motion, with little dependence on the precise composition of the stimulus or its spatial position. Thus the challenge of reducing the spatial specificity of perceptual learning can be formulated in terms of increasing the contributions of neurons in higher-level cortical areas to the perceptual response (Dosher, Jeter, Liu, & Lu, 2013).

Previous work using causal manipulation of neural activity has shown that the perceptual reweighting of visual areas is highly sensitive to the training procedure. Specifically, the stimulus used during training has a profound effect on the contribution of individual brain structures, as assessed by protocols in which specific areas are inactivated at different times relative to the training (Chang, Mevorach, Kourtzi, & Welchman, 2014; Chen, Cai, Zhou, Thompson, & Fang, 2016; Chowdhury & DeAngelis, 2008; Liu & Pack, 2017; Walsh, Ashbridge, & Cowey, 1998). These results show that greater weight can be accorded to different brain structures by training with appropriate stimuli.

These considerations lead to a straightforward hypothesis: Training with stimuli that are selectively encoded in higher-level structures will lead to perceptual learning with less specificity (Das, Tadin, & Huxlin, 2014; Fine & Jacobs, 2002; McGovern, Webb, & Peirce, 2012; Zhang & Tadin, 2019), both for spatial position and for stimulus features. Recent evidence in support of this idea comes from a study (Liu & Pack, 2017) in which the contribution of the middle temporal (MT) area to motion perception in non-human primates was found to depend on the stimulus used during training. Inactivation of MT devastated performance after the animals had been trained on complex random dot kinematograms, but had little effect when they had been trained on simple grating stimuli. The effects of training with complex stimuli transferred to simpler stimuli and showed some evidence of greater spatial generalization.

In this study, we examine the relationship between stimulus complexity and the specificity of perceptual learning. Consistent with previous work (Fahle, 1997; Huxlin et al., 2009), we find that observers trained to discriminate the properties of grating stimuli exhibit specificity for spatial position. In contrast, observers trained with more complex stimuli, such as optic flow patterns, exhibit far less specificity. Reduced spatial specificity is associated in all cases with greater spatial integration, lending strong support for the idea that training with complex stimuli leads to a reweighting of the perceptual contributions of higher level cortical areas.

## Methods

### Experimental Procedure

#### Observers and Apparatus

All observers were naïve to visual psychophysics and unaware of the purpose of the study. Each observer gave written, informed consent prior to participation in the study, which was approved by the Ethics Committee of the Montreal Neurological Institute and Hospital (NEU-06-033).

The stimuli were generated in MATLAB with Psychtoolbox (Brainard, 1997), and presented on a 21-inch *hp* Trinitron CRT monitor (1024 pixel × 768 pixel, 0.37 mm [H], 0.37 mm [V] per pixel, 85 Hz frame rate). Observers were seated 57 cm from the screen, and their heads were stabilized with a chin-rest. Experiments were run in a normally lit room.

#### Visual Stimuli

##### Experiment 1

This experiment had two training phases: (1) drifting grating (DG) motion discrimination (left vs right), and (2) translation random-dot kinematogram (RDK) motion discrimination (left vs right). The DG stimuli were Gabor patches, with spatial and temporal frequencies set to 1 cycle/degree and 8 cycles/second. The RDK stimuli consisted of small (0.1°) white and black dots, at a density of 2.6 dots/deg.^2^, presented on a grey background (luminance 63.1 *cd/m*^2^). Each stimulus was windowed inside a circular aperture 6 degrees in diameter. The dots presented on each frame either belonged to the signal or to the noise group. The signal dots moved coherently in a specific direction, while the noise dots moved randomly in different directions. The coherence level of the random-dot stimulus determined the proportion of the dots that belonged to the signal group. The horizontal and vertical velocity of each dot followed the equation below:

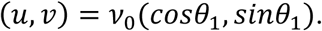

For dots that carried the global motion signal, *θ*_1_was set to 0 for leftward motion and *π* for rightward motion. For the noise dots, the value of *θ*_1_ was chosen randomly between (0,2*π*). *x* and *y* denote the horizontal and vertical location of each dot in the visual field (in degrees). *v*_0_ represents the dot velocity and was set to 10°*s*^−1^. For both the RDK and DG stimuli, the duration on every trial was 70.6 ms, corresponding to 6 frames at the 85 Hz refresh rate. The stimulus during training was positioned in the upper fight visual field at an eccentricity of 5 degrees.

##### Experiment 2

This experiment had two training phases: (1) translation RDK motion discrimination (left vs right), and (2) optic flow random-dot motion discrimination (expansion vs contraction). The translation RDK stimulus was identical to the one used in the second phase of Experiment 1. For the optic flow stimulus, the horizontal and vertical velocity of each dot followed the equation below:

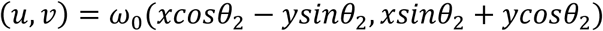

For signal dots, the value of *θ*_2_ was set to *π*/2 or 3*π*/2 for expansion or contraction, respectively. For the noise dots, the value of *θ*_2_ was chosen randomly from (0,2*π*). We set the radial velocity *ω*_0_ to 4 Hz. Coherence was modulated in fashion analogous to that used for the translation RDKs.

While all observers were able to perform the translation RDK task well on the first session, we noticed in pilot experiments that some observers initially struggled with the optic flow stimulus. Given the longer temporal integration for optic flow compared to translation RDK (Burr & Santoro, 2001), we therefore used a slight longer duration for these stimuli (94.1 ms, corresponding to 8 monitor frames).

#### Training Procedure

Every observer completed 8-10 days of training sessions for each phase of the two experiments. In each training session, observers completed 5 blocks of training, where each block consisted of 125 trials that lasted in total about 5-6 minutes. Between the training blocks, observers took a 1-minute break, after which the next block automatically began. In total, each training session was about 30 minutes long.

At the beginning of each training block, the stimulus coherence in the random-dot stimuli was set to 0.7, and the stimulus contrast in the DG stimulus was set to 0.5. Both coherence and contrast changed in an adaptive procedure (see below) to maintain the difficulty level throughout the training blocks and sessions for the two tasks.

Each trial began with the acquisition of fixation on a central point. Eye position was monitored throughout each trial (EyeLink 1000) and required to be within 1 degree of the fixation point for the trial to be included in subsequent analyses. After each stimulus presentation, observers reported their perception using a keyboard button press, which then started the next trial. Observers were compensated for their time in proportion to the number of correct responses (¢1 per correct trial).

#### Testing Procedure

After each training phase, observers were tested on spatial integration and specificity. The stimulus used for the testing procedures at the end of both training phases was the simpler stimulus: DGs for Experiment 1, and translation RDK motion for Experiment 2.

For the spatial integration task, the stimulus was presented in the same location as the training stimulus, but the discrimination threshold was measured at different stimulus sizes. The threshold for DG contrast was evaluated at stimulus diameters of 1.25, 5, 8.75, 12.5, 16.25, and 20 degrees. The threshold for translation RDK motion discrimination was evaluated at diameters of 6, 12, 18, 24, 30, and 36 degrees.

To measure the spatial specificity of the learned improvement, the test stimulus was presented with the same aperture size as in the training, but in different locations of the visual field. The eccentricity of the stimulus was preserved for different test locations (5 degrees). Eight positions on the vertical, horizontal, and oblique meridians of the visual field were used for the transfer evaluation (Figure 2b).

**Figure 2:**
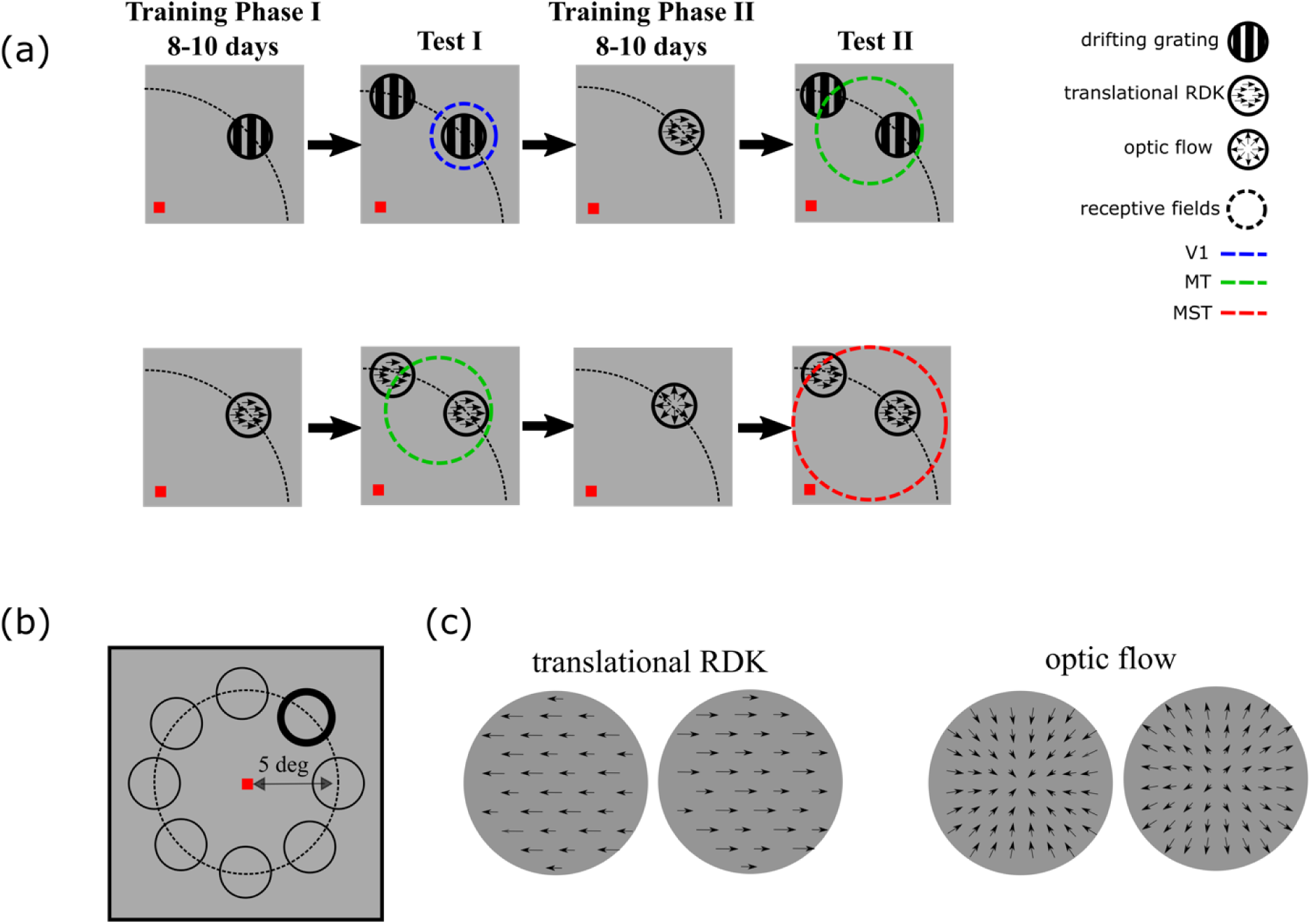
The experimental paradigm. (a) Schematic presentation of our experimental paradigm: Each square represents the upper-right quadrant of the visual field. The small, red square (at the lower-left corner of each square) shows the central fixation point. Training on drifting grating for 8-10 days (top row, first column) increases the readout weight of an area with small receptive fields, such as V1 (top row, second column). Because of the small receptive fields (blue dashed circle), learning will not generalize to stimuli at different locations. Training on translation RDK stimuli for 8-10 days (top row, third column) instead increases the readout weight of an area with larger receptive fields, such as MT (top row, fourth column). Because of the larger receptive fields (green dashed circle), learning will generalize to more distant locations. The same argument suggests that training on a complex optic flow stimulus (bottom row, third column) increases the readout weight of an area with even larger receptive fields, such as MST. Spatial specificity will be lower, as the recruited receptive fields of MST neurons are larger (bottom row, fourth column). (b) The configuration of the trained (thick black circle) and tested (thin black circles) stimulus locations that was used in Experiments 1 and 2. The circles show the stimulus locations, and all had the same distance (5 degrees) from the center of the screen (red square). (c) Example vector fields for translation RDK (left) and optic flow (right). The direction and size of each vector represents the motion direction and velocity of a moving dot.

### Staircase procedure

A standard 2-down-1-up staircase procedure was used for setting the coherence of the random-dot stimulus (Leek, 2001). The staircase procedure resulted in an 83% convergence level. The last six reversals of the staircase were used to estimate the discrimination thresholds.

### Statistical Analysis

The discrimination threshold was measured using the staircase procedure in each training block. The thresholds were then averaged over training blocks on each day to track the performance improvement across days.

As explained above, to evaluate the spatial integration after each training phase, observers’ discrimination thresholds were measured with different stimulus sizes. To obtain a quantitative measure of spatial integration, we fitted a difference of error functions to the threshold values as a function of stimulus sizes (Pack, Hunter, & Born, 2005; Uka & DeAngelis, 2003):

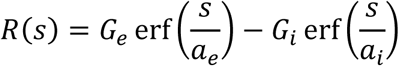

*R*(*s*) represents the discrimination threshold at stimulus size *s*. In this model, the threshold at every stimulus size is determined by an excitatory 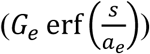) and an inhibitory 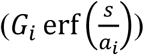 components that interact through subtraction. The gains of the excitatory and inhibitory components (*G_e_* and *G_i_*), and the excitatory and inhibitory area sizes (*a_e_* and *a_i_*) were estimated by fitting the model to the measured thresholds. We used a nonlinear least square curve fitting method to fit the model to our data (*lsqcurvefit* MATLAB function with Trust-Region-Reflective optimization). The estimated excitatory area *a_e_* was used as the spatial integration index (SI).

To quantify the spatial specificity of the learning effect, we fitted a linear model to the measured thresholds as a function of distance from the training location. The slope of the fitted line was used as the specificity index. Higher specificity leads to larger slopes, since the threshold increases as the distance from the trained location increases. Slope values that are close to zero show threshold values that are almost equal for different distances from the trained location.

We used linear mixed-effect analysis for statistical comparisons between conditions. The experimental conditions (e.g. post-training phase 1) were used as categorical fixed-effect variables, and the measured spatial integration or spatial specificity were set as the output variables. The identity of the observers was used as the random-effect. We used the *fitlme* function in MATLAB to fit the mixed-effect models to our data.

### Computational model

Our computational model consisted of a sensory visual representation stage that encodes the visual stimuli, and a decoder that reads out from the sensory neurons to make a perceptual decision (Figure 6a). The sensory representations are assumed to be located in area MT, which is known to encode translation motion of the kind we studied in Experiment 1 (Cui, Liu, Khawaja, Pack, & Butts, 2013; Dubner & Zeki, 1971; Lagae, Maes, Raiguel, Xiao, & Orban, 1994) and area MST, which encodes the optic flow stimuli that we studied in Experiment 2 (Duffy, 1998; Lagae et al., 1994; Mineault, Khawaja, Butts, & Pack, 2012). The decoder is simply a linear readout that could be implemented in many different brain regions.

**Figure 6:**
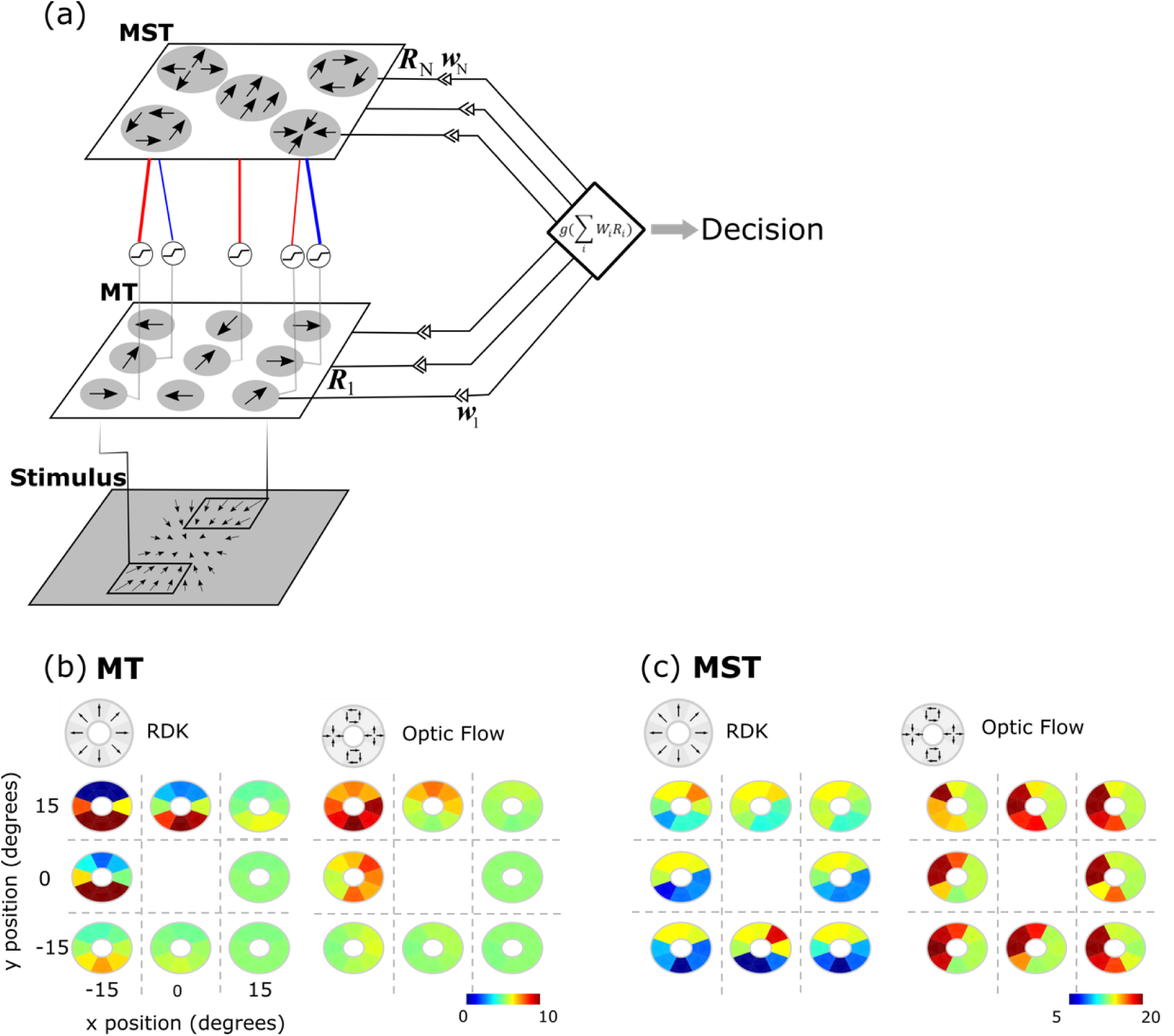
The structure of our computational model and example simulated MT/MST neurons. (a) Structure of our computational model: the stimulus is first processed by a population of MT-like neurons that decompose the stimulus into a distributed representation of motion directions and locations. MT neurons project to the next motion processing stage that forms the MST stage of our model. The synaptic connections between MT and MST neurons can be inhibitory (in blue) or excitatory. Before projecting to MST, the output of each MT neuron passes through a nonlinearity. The output of each MT and MST neuron (*R_i_*) is sent to an adaptive decoder for linearly decoding the motion direction. (b) Tuning curves of an example MT neuron: tuning mosaics demonstrate the firing rate of the neuron to every motion type (RDK in left and optic flow in right). The tunings are measured in eight different positions around the visual field (every mosaic corresponds to a spatial position). (c) Similar to (b) but for an example MST neuron.

The structure of our MT-MST model followed a previously established hierarchical model of the dorsal visual pathway (Mineault et al., 2012) (Figure 6a). Model MT neurons had retinotopic two-dimensional Gaussian receptive fields that spanned the visual field (−15° *to* 15° horizontal and vertical visual angles at 0.2° resolution):

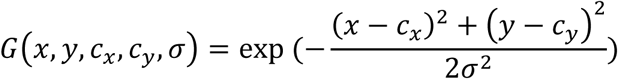

Where *c_x_* and *c_y_* are the horizontal and vertical coordinates of the receptive field center, and *σ σ* is the width of the receptive field set to be 2 or 4 degrees. The MT neurons were also tuned to different motion directions and speeds, which were modeled as follows:

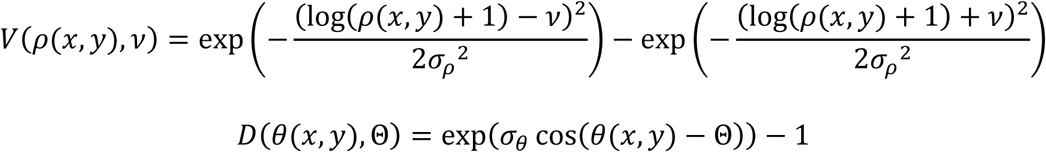

where, preferred speeds (*v*) and direction (Θ) are chosen from {5, 12, 30} 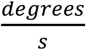 and {0, 45, 90, 135, 180} *degrees*, respectively. The speed tuning width *σ_p_* was set to *σ_p_* = 1, and the direction tuning width was chosen to be *σ_θ_* = 2.5 similar to the tuning properties of MT neurons reported in MT (Albright, 1984; Nover, Anderson, & DeAngelis, 2005).

The direction, speed, and position tunings were then combined to determine the firing rate of the neuron:

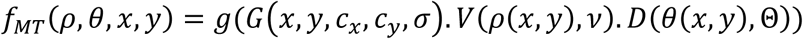

where *g* = max(*x*, 0)^0.2^ is a compressive nonlinearity that has been shown to be essential in building MST-like tuning from MT responses (Mineault et al., 2012).

Every MST unit received inputs from *N* MT neurons (*N* = 15 here) with receptive fields centered within the receptive field of the MST unit. The *N* MT neurons were linearly combined:

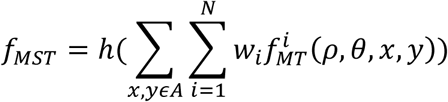

where *w_i_* is the synaptic strength between the *i^th^* MT neuron and the MST neuron, which can be excitatory (positive values) or inhibitory (negative values). *w_i_* was sampled randomly from (0,1] or [−1,0), with 0.2 probability of being sampled from the negative range. *A* represents the receptive field of the MST neuron. ℎ = max (*x*, 0) is a half-wave rectification nonlinearity. *f_MST_* was spatially convolved with the output of the MT representation of the stimulus, and then spatially subsampled (max-pooling) to form the MST output (Fukushima, 1988). The combination of MT inputs with different direction, speed, and position tunings rendered MST neurons sensitive to more complex motion patterns, while the spatial convolution and max-pooling helped to decrease the spatial specificity of the representation at the MST stage compared to MT (Fukushima, 1988).

The output of the MT and MST population were then passed to an adaptive decoder that linearly combined the responses of the neurons and decided on the motion direction. The adaptive decoder has been previously proposed to model different aspects of visual perceptual learning in humans (Jacobs, 2009). The decoder linearly decodes motion direction given the output of a subset of sensory neurons. Initially, the adaptive decoder pools the output of M randomly chosen sensory neurons (across MT and MST population; here M = 5) to determine the probability of one motion direction versus the other:

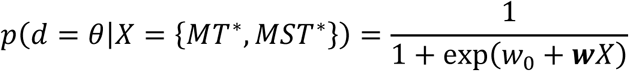

Throughout the training, in every iteration, the adaptive decoder drops the least informative neuron, and randomly selects another neuron to add to the pool of readout neurons. After many iterations, the decoder converges to a subset of sensory neurons that are most optimal for solving the training task. The discrimination threshold of the decoder is determined by finding the stimulus coherence or contrast that yields 85% decision accuracy.

## Results

Our experiments are motivated by physiological observations in the dorsal visual pathway, which is specialized for motion processing. As shown in Figure 1, neuronal receptive field sizes increase as one follows the dorsal pathway from V1 to MT to MST (Gattass & Gross, 1981; Raiguel et al., 1997). Crucially for our study, the increase in receptive field sizes is accompanied by changes in the stimulus selectivity in each area. Specifically, tuning for simple motion stimuli, such as drifting gratings, is found at every stage along the pathway, though it decreases at higher-level stages (Khawaja, Liu, & Pack, 2013). In contrast, preferences for more complex motion stimuli, such as optic flow, emerge in MST (Duffy & Wurtz, 1991; Lagae et al., 1994; Tanaka, Fukada, & Saito, 1989).

**Figure 1:**
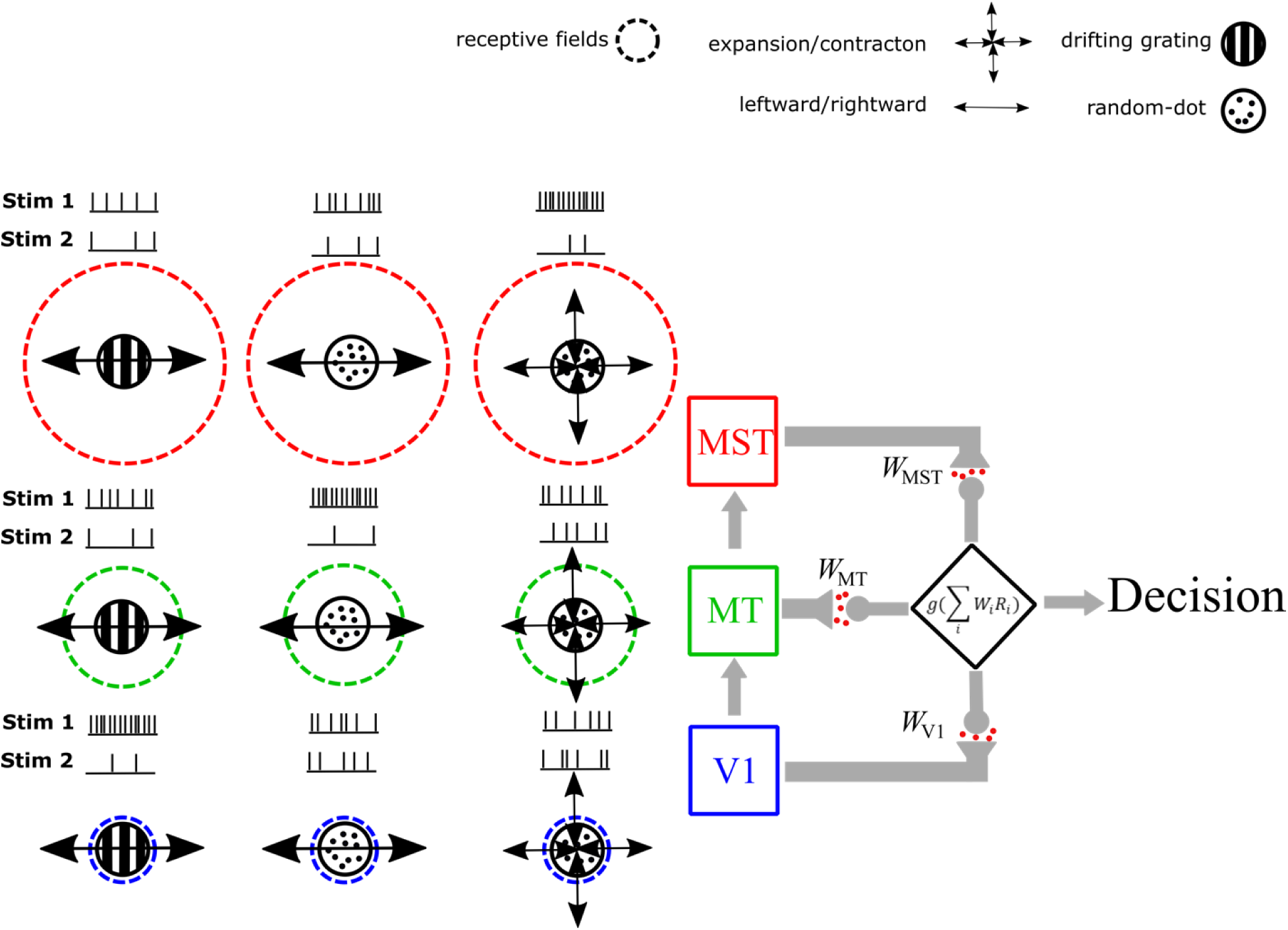
Schematic presentation of the dorsal visual pathway: Areas V1, MT, and MST are three visual areas that encode motion stimuli in the dorsal pathway. Their stimulus selectivity is different as higher areas prefer more complex moving patterns. The small vertical lines next to each area represent the firing rates of a typical neuron in each area evoked by each stimulus type. Stimulus 1 and 2 represent the preferred and null motion direction, respectively, for a typical neuron in each area (i.e. leftward/rightward motion for DG and RDK, or expansion/contraction for optic flow). The size of the receptive fields also increase from lower to higher areas. Colored dashed circles show the ordinal arrangement of receptive field size of each area (not drawn to scale). Each sensory visual area projects to a downstream sensorimotor/decision making area that integrates inputs from the three areas. The contribution of each sensory area to the decision or action depends on it readout weight (*W*_*v*1_, *W_MT_*, *W_MST_*). According to our hypothesis, training with the stimulus that elicits the highest selectivity in an area (e.g. drifting grating in V1) increases the readout weight of that area (e.g. *W*_*v*1_).

Given the standard assumption that perceptual learning entails a reweighting of the contributions from neurons in these different areas (Dosher et al., 2013), we reasoned that the spatial properties of perceptual learning should be influenced by the complexity of the stimuli used during training. Specifically, spatial integration – the propensity of observers to combine visual signals across spatial locations – should increase following training with complex stimuli, as the contribution of neurons with larger receptive fields is increased. For the same reason, the spatial specificity of learning should be lower after training with more complex stimuli, since neurons with large receptive fields generalize stimulus selectivity across space. In this section, we test these hypotheses with two psychophysical experiments, and then examine a computational model that formalizes the intuitions illustrated in Figure 1.

### Experiment 1: drifting grating vs random dots

Our first set of experiments made use of drifting grating (DG) stimuli (Gabor patches) and random-dot kinematograms (RDKs). DG stimuli are well-matched to the receptive field properties of orientation selective neurons in the primary visual cortex (V1) and other low-level cortical areas. They also contain motion information that is available locally, with no need for spatial integration. In contrast, RDKs are complex, because they are noisy and hence accurate estimation of their motion requires integration across space and local motion directions (Britten, Shadlen, Newsome, & Movshon, 1993). As a result, these stimuli yield poor selectivity in areas like V1 (Snowden, Treue, & Andersen, 1992), but strong selectivity in higher-level cortical structures, such as the middle temporal (MT) (Albright, 1984; Britten et al., 1993) and medial superior temporal (MST) (Duffy, 1998) regions.

In this and subsequent experiments, we probed the spatial properties of perceptual learning in two ways. The first measure was spatial integration, defined as the task performance for stimuli of different sizes. This provides a measure of the extent to which observers integrate across space, and it correlates with receptive field sizes in the brain regions contributing to perceptual decisions (Liu, Haefner, & Pack, 2016; Tadin, Silvanto, Pascual-Leone, & Battelli, 2011). The second was a direct measure of spatial specificity: We tested psychophysical performance for stimuli placed at different positions in the visual field. According to the logic outlined in the Introduction, training with simple stimuli, such as DGs, should yield high specificity with little spatial integration, while training with RDKs should decrease specificity and increase integration (Figure 2a).

Six observers were recruited for this experiment and underwent two phases of perceptual training. In the first phase (8-10 days), they were trained to discriminate between leftward and rightward motion of the DG stimuli. The contrast of the grating was modulated in a staircase procedure on every trial (see Methods). In the second phase (8-10 days), the same observers were trained on leftward/rightward motion discrimination with RDKs. The coherence of the moving dots was modulated on every trial in a staircase procedure. The training stimuli in both phases were of the same size (6 degrees in diameter). At the end of each phase, each observer’s motion discrimination performance across stimulus sizes was measured with the DG stimulus. This was done to provide a fair comparison across training phases, but as we show below, the results are not specific to the DG stimulus. Performance in this and in subsequent experiments was defined in terms of sensitivity, which is the reciprocal of the threshold contrast or coherence.

### Spatial Integration

As shown in Figure 3a, the motion sensitivity of observers improved in both training tasks (*first phase*: *F*_(1,10)_ = 7, *p* = 0.024, – *second phase*: *F*_(1,10)_ =22.47, *p* < 0.001), demonstrating the effectiveness of the training. As outlined in the Introduction, if training with complex stimuli leads to an increase in the weighting of neurons in higher-level cortex, observers should exhibit increased spatial integration after training with RDKs. That is, their motion discrimination performance should improve for larger stimuli after training with complex stimuli.

**Figure 3:**
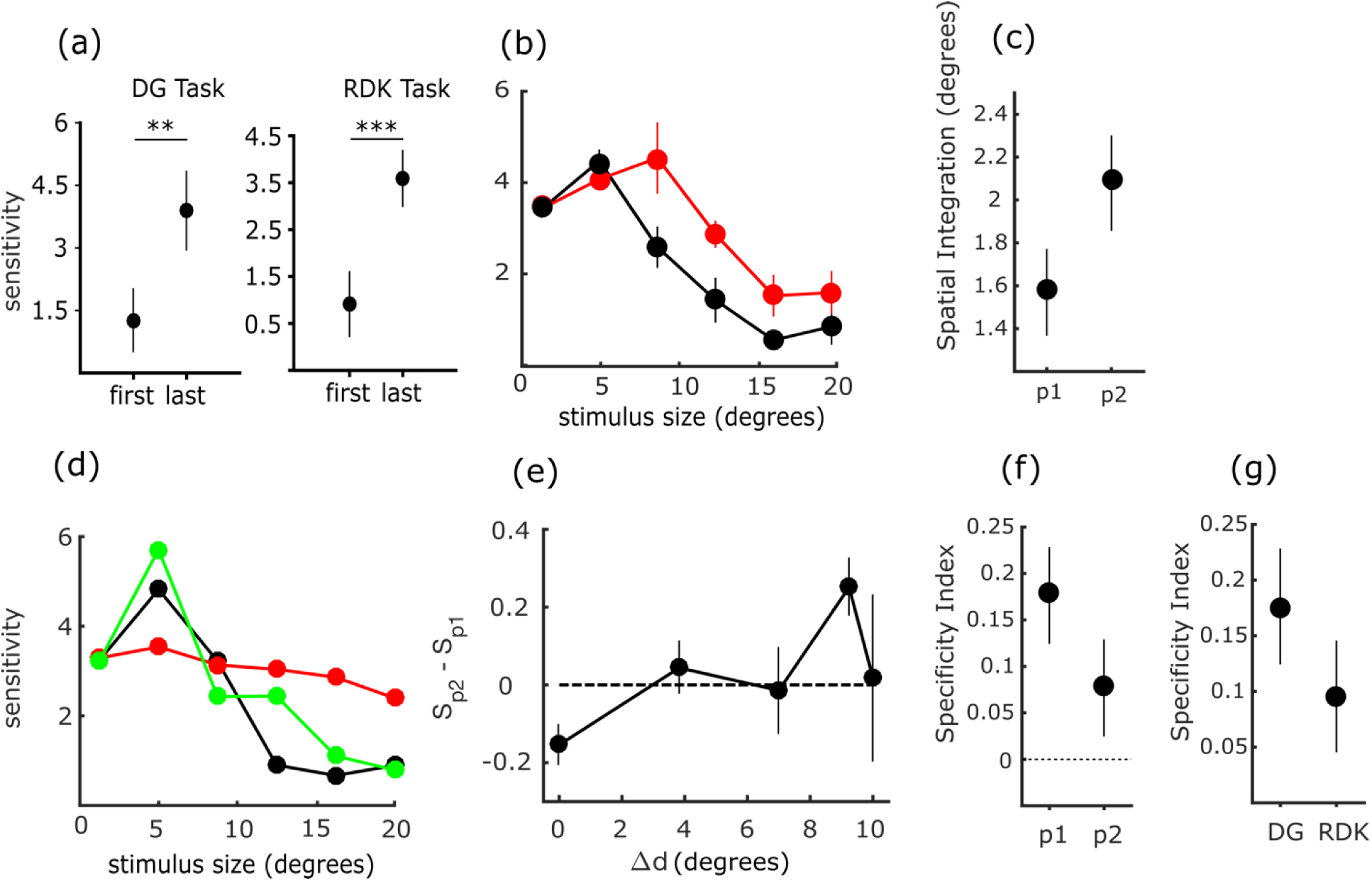
(a) Change in sensitivities in first phase (DG) and second phase (translation RDK). (b) Log(sensitivity) for different stimulus sizes averaged across observers – post phase 1 (DG in black) and post phase 2 (RDK in red). (c) Spatial Integration measured with DG for post-DG and post-RDK. (d) Log(sensitivity) for different stimulus sizes for an example observer – post phase 1 (DG in black), post phase 2 (RDK in red), and post phase 3 (DG in green). (e) Changes in Log sensitivity for different distances from trained locate--on measured with DG from the first to the second training phases (*S_p2_* − *S_p1_*). (f) Specificity index tested with DG after phase 1 (post DG) and after phase 2 (post RDK). (g) Specificity index after training on DG and tested on DG (DG) and after training on RDK and tested on RDK (DG). Error bars show SEM.

Figure 3b shows the spatial integration curve measured at the end of the first and the second training phases (averaged across observers). After training with DGs, the spatial integration curve (black line) peaked for a stimulus size of approximately 5 degrees and then decreased sharply with increasing stimulus size, as reported previously (Tadin et. al., 2003). After training with RDKs, we again tested spatial integration with DGs and found that the peak of the spatial integration curve (red line) moved rightward, so that the best performance was obtained for a stimulus size of approximately 10 degrees. This result cannot be attributed to increased familiarity with the task or to a general improvement in motion processing, because the observer’s performance actually decreased slightly for smaller stimuli.

To quantify these effects for the population of observers, we fitted a parametric model to the spatial integration curves of every observer to estimate their spatial integration size (see Methods). All six of the observers who participated in this experiment showed an increase in spatial integration between the first and the second training phases (*F*_(1,10)_ = 8.08, *p* = 0.017) (Figure 3c).

### Spatial Specificity

Most previous studies have found that improvements in performance after perceptual learning are specific to the trained location, although recent work has demonstrated that specificity can be decreased with appropriate training protocols (Watanabe & Sasaki, 2015). From a theoretical perspective, the degree of spatial specificity is likely to be associated with the size of the receptive fields of the neurons contributing to the perceptual task (Dosher et al., 2013). We therefore predicted that training protocols that increase spatial integration, as in our results with RDKs (Figure 3c), should also exhibit less spatial specificity (Figure 2a). To test this idea, at the end of each training phase, we measured sensitivity across different spatial locations with DG stimuli (Figure 2b). To avoid confounds due to overall visual sensitivity, we examined performance at locations with the same retinal eccentricity as the trained location (thick black circle) but arranged symmetrically around the visual field. Performance was measured after each phase of the experiment.

Figure 3e shows the change in sensitivity for DGs between phase 1 (*S_p1_*) and phase 2 (*S_p2_*), as a function of distance from the training location for the group of observers. Here distance (Δd) is expressed as the distance from the center of the stimulus at the training location to the center of each test stimulus (Figure 2b). After RDK training, the sensitivity of DG motion discrimination improved at locations distant from the trained location. The improvement was greatest at a distance (Δd) of 9.2 degrees of visual angle from the training location (*p* = 0.0137), indicating that training with RDKs extended to different locations and different stimuli. This improvement in performance was not due to repeated exposure to the task locations, because sensitivity actually decreased at the trained location (*p* = 0.0147). We provide an explanation for this counterintuitive result below.

To quantify the specificity of perceptual learning across the training protocol, we defined a specificity index (Methods), which takes on a value of zero when sensitivity is equal at all spatial locations. Positive values indicate higher sensitivity for the trained location. Specificity was high after the first phase of training (mean specificity index = 0.1763), consistent with previous reports showing a lack of transfer of perceptual learning across spatial positions for Gabor stimuli (Fahle, 2005). However, after the second phase of training, the specificity index decreased significantly to 0.076 (*phase*1: 0.1763 ± 0.052 *vs*. *phase*2: 0.076 ± 0.0524) (Figure 3f-left). Thus, the level of performance obtained after training with RDKs was largely maintained when the stimulus was moved or changed.

One limitation of these results is that the observers were tested twice on the spatial specificity task with the DG stimuli. Thus, some improvement after phase 2 could be attributed to prior exposure to the stimulus at the untrained location, rather than to the properties of the training stimulus. Likewise, our protocol could be considered an instance of “double-training” (Xiao et al., 2008), which has been shown to decrease specificity. To examine these possibilities, we trained a second group of nine observers *only* on the RDK task and measured their spatial specificity for the same RDK stimulus. We then compared this specificity to that measured in the first group, after the first phase of training and testing with DGs. Thus, in this comparison, neither group had prior exposure to the stimuli at the testing locations; the only difference was in the stimuli to which the two groups were exposed. Moreover, each group had only participated in a single phase of training at the time that they were tested for specificity. As shown in Figure 3f (right), the group that was trained on RDKs exhibited less spatial specificity compared to the group trained on the DGs (*Wilcoxon ranksum test*; *p* < 0.01), suggesting that the structure of the training stimulus has an important effect on spatial specificity, apart from other aspects of the task design (Watanabe et al., 2002).

### Task Specificity

An alternative possibility is that the increased spatial integration after RDK training reflects a more general strategy of integrating out noise, irrespective of the specific structure of the signal. In that case, training on any task that requires filtering out noise would increase behavioral integration. To test this idea, we trained seven observers on a task that involved similar noise levels as the RDK task, but did not require motion discrimination. Specifically, we trained observers on a face discrimination task (8-10 days), in which two different faces were shown at the beginning of every training block (face A and face B), and observers were asked on every trial whether a probe face was A or B. Spatial noise was added to the stimuli at levels determined by a staircase procedure (see Methods). As the face discrimination task did not require motion processing, any change in spatial integration of motion stimuli could be attributed to the spatial noise in the training stimulus.

We measured spatial integration of DG motion stimuli before and after training on face discrimination. Across observers, there was no consistent change in the spatial integration of motion signals after the face discrimination training (*F*_(1,12)_ = 0.712, *p* = 0.42). This observation shows that the learning effects after RDK training were not due to a general improvement in the ability to filter out spatial noise. From a physiological perspective, this suggests that learning does not transfer from the ventral to the dorsal stream.

### Experiment 2: simple translation motion vs complex radial motion

The results thus far can be interpreted in terms of the function of the dorsal visual pathway. Early stages of this pathway encode the motion of DG stimuli with high fidelity (Swindale, Matsubara, & Cynader, 1987; Weliky, Bosking, & Fitzpatrick, 1996), while robust encoding of the motion of RDK stimuli emerges in later stages, notably area MT. We argue that the larger receptive fields of neurons in the later stages accounts for many of the results seen in Experiment 1.

Continuing this line of reasoning, we next consider the terminal stage of the dorsal pathway, area MST. Compared to neurons in MT, MST neurons have far larger receptive fields (Duffy & Wurtz, 1991) and encode more complex motion patterns, such as radial and circular motion (Cui et al., 2013; Duffy & Wurtz, 1991; Lagae et al., 1994; Mineault et al., 2012). We refer to these latter patterns as optic flow (Gibson, 1950). Quantitative studies of neuronal populations have shown that translation motion of the kind used in Experiment 1 is encoded more effectively in MT than in MST, while the encoding of radial and circular motion patterns is more robust in MST than in MT (Cui et al., 2013; Lagae et al., 1994; Mineault et al., 2012). We therefore predicted that perceptual training with optic flow should further increase the readout weight of MST neurons, which should in turn manifest behaviorally as increased spatial integration and decreased specificity.

As in the previous experiment, observers underwent two phases of training, the first with a relatively simple motion stimulus and the second with a more complex stimulus. To facilitate comparison of training effects, we again probed spatial integration and specificity with the simpler stimulus set.

Nine observers were recruited for this experiment. Figure 2a (bottom) shows a schematic of the experimental paradigm. Observers went through two phases of perceptual training. In the first phase (8-10 days), they were trained to discriminate the leftward/rightward motion of the RDK stimulus described above (These data were mentioned above in the presentation of Experiment 1 (Figure 3f)). In the second phase (8-10 days), the same group of observers were trained to discriminate between expansion and contraction optic flow, also defined with noisy random-dot patches. The latter requires integration across motion directions to estimate the pattern (Figure 2c). Importantly, the sizes of the training stimuli were again identical across the two phases of training.

#### Spatial Integration

Observers improved during both training tasks (Figure 4a) (*First phase*: *F*_(1,16)_ = 30.866, *p* < 0.001 − *second phase*: *F*_(1,16)_ = 23.94, *p* < 0.001), indicating that the training protocol was effective. Figure 4b shows the spatial integration measured after phase 1 (black curve – averaged across observers). The spatial integration peaked at around 10 degrees. After training on optic flow motion (red curve in Figure 4b), however, motion sensitivity for the same translation stimulus increased monotonically with stimulus size, indicating much stronger spatial integration. Moreover, the sensitivity at smaller stimuli actually decreased after optic flow training (compare red and black curves at 4- and 10-degree stimulus sizes). Similarly, for the group of observers, the mean spatial integration (Figure 4c) increased sharply after training on complex motion (*F*_(1,16)_ = 27.1, *p* < 0.001).

**Figure 4:**
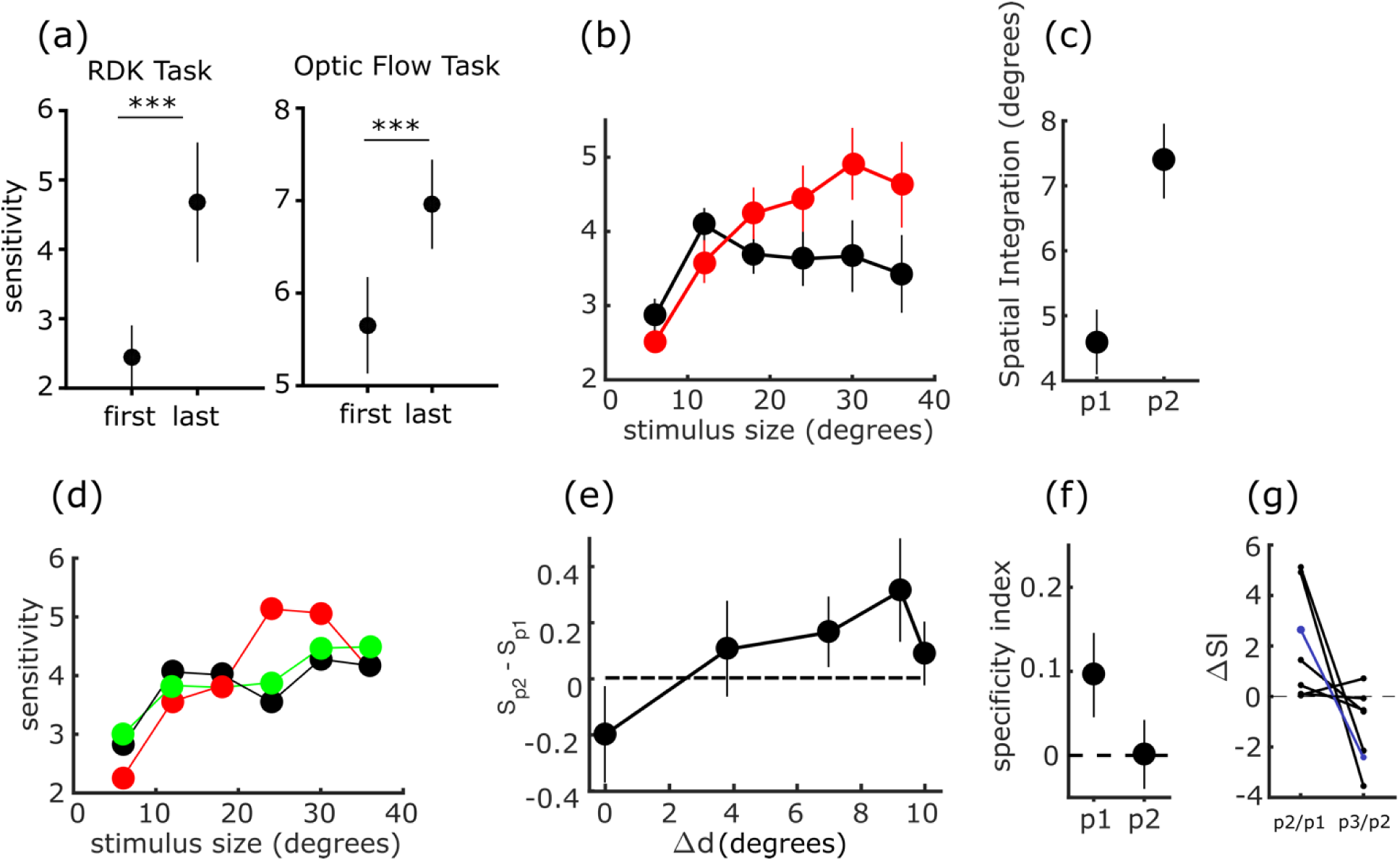
(a) Change in sensitivities in first phase (translation RDK) and second phase (radial RDK). (b) Sensitivity for different stimulus sizes for the average of observers – post phase 1 (translation RDK in black), and post phase 2 (radial optic flow in red) (c) Spatial Integration measured with translation RDK for post-RDK and post-optic flow. (d) Sensitivity for different stimulus sizes for an example observer post phase 1 (RDK in black), post phase 2 (optic flow in red), and post phase 3 (RDK in green) (e) Changes in sensitivity at different spatial distances from the trained location measured with (*S_p2_* − *S_p1_*). (f) Specificity index tested with RDK after phase 1 (post RDK) and after phase 2 (post optic flow). (g) The changes in spatial integration (Δ*SI*) are shown for observers who went through three phases of training. For all observers, spatial integration increased from the end of the first phase to the end of the second phase (positive values for p2/p1). 6 out of 7 observers showed a decrease in the spatial integration from the end of the second phase to the end of the third phase (negative values for p3/p2). The black lines and circles show the changes in spatial integration for individual observers. The blue line shows the change in spatial integration for the observer shown in (b). Error bars show SEM.

#### Spatial specificity

As in the previous Experiment (Figure 3e), observers were tested with the simpler stimuli (translation motion in this case) after completion of each phase of training at a range of positions having equal eccentricity but different angular locations relative to the trained location (Figure 2b). Figure 4e shows the change in sensitivity for translation motion between phase 1 and phase 2 as a function of distance from the training location (*S_p2_* − *S_p1_*) for the group of observers. Training on optic flow improved sensitivity to translation motion at locations distant from the trained location (at Δ*d* = 9.2°, *F*_(1,8)_ = 8.1747, *p* = 0.017, Δ*d* = 7°, *F*_(1,8)_ = 5.8764, *p* = 0.041), but decreased the sensitivity substantially at the trained location (*F*_(1,8)_ = 10.875, *p* = 0.010) (Figure 4e). For the group of observers, the specificity index decreased significantly after the second training phase (*phase*1: − 0.095 ± 0.05 *vs*. *phase*2: 0.0012 ± 0.041) (Figure 4f).

Overall, the results of this experiment support the idea that more complex optic flow stimuli can increase the readout weight of neurons in higher areas of the dorsal pathway that are tuned to more complex moving patterns, such as MST.

#### Reversal of training effects

As mentioned above, these training effects were unlikely to be a simple consequence of double-training, since they can be observed after a single round of training in intergroup comparisons (Figure 3f). Nevertheless, we sought to ensure that our results could not be attributed to the fixed sequence of training phases (simple stimuli followed by complex stimuli) in our design.

Following the completion of Experiments 1 and 2, we were able to retain a subgroup of 7 observers for a third phase (3 for Experiment 1 and 4 for Experiment 2). In this third phase, we retrained the observers on the same stimulus to which they had been exposed on the first phase, with training lasting another 8 days. This experiment served as a further control against the possibility that the increases in spatial integration seen in both experiments were due to cumulative exposure to the motion task. If this were true, spatial integration should continue to increase after the third phase.

Data from an example observer in Experiment 1 are shown in Figure 3d. This observer’s spatial integration curve had shifted rightward following the second phase of training (black and red curves after phase 1 and 2 respectively), but after the third phase it shifted leftward (green curve in Figure 3d), as predicted from our main hypothesis. A similar finding was obtained with an example observer from Experiment 2, as shown in Figure 4d (green line).

All 7 of the observers who completed a third training phase had shown an increase in spatial integration at the end of the second phase compared to the first phase (Figure 4g; *F*_(1,12)_ = 36.322, *p* < 0.001). After the third training phase, in the majority of observers (6 out 7), the spatial integration decreased compared to the end of the second training phase (*F*_(1,12)_ = 5.44, *p* = 0.0378). These results suggest that increases in integration do not follow from repeated task performance but rather depend on the stimuli used during training.

#### Relationship between learning depth and change in spatial integration

The changes in spatial processing that we observed in the previous experiments could be a result of learning in the first phase, the second phase, or both. In other words, it could be that training on the first phase of the experiment actually decreases spatial integration and increases spatial specificity relative to the baseline, and that the second task simply undoes the effect of the first phase. Or the second phase of training could increase spatial integration and decrease specificity by a reweighting mechanism that is independent of the effects of the first training phase. Both strategies would show the same outcome in terms of change in the spatial processing from the first to the second phases.

We reasoned that, to the extent that each training phase is responsible for the reweighting of neural inputs to perceptual decisions, the depth of learning observed during a given phase of training should predict the subsequent changes in integration and specificity. We therefore quantified the depth of learning in both training phases for every observer (Δ*T*_1_, Δ*T*_2_), as well as the change in the spatial integration or specificity from the first phase to the second phase (Δ*S*). Then, we fit a linear model to predict the observed changes based on the improvements at the trained location in the first and the second phases of both experiments:

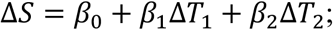

The estimated *β_i_* value indicates the variance in Δ*S* across observers that can be explained by depth of their learning in the *i*^*th*^ training phase.

We pooled the data from Experiment 1 and Experiment 2 to fit the linear regression model. Our fitted model (*R*^2^ = 0.62) showed that the improvement in performance during the first training phase failed to predict subsequent spatial integration (*p* = 0.43). In contrast, learning during the second phase exhibited a positive, statistically significant effect (*p* < 0.01). The estimated *β* coefficients for the two training phases are shown in Figure 5a (left). Applying a similar linear regression model to the measured changes in spatial specificities in Experiments 1 and 2 yielded a similar result (*R*^2^ = 0.54): The changes in the spatial specificity across observers were predicted by the depth of learning in the second training phase (*p* = 0.019), and again the first training phase had a negligible effect (*p* = 0.24). Figure 5a (right) shows the estimated *β* coefficients for the two training phases. This result supports the idea that the changes we observed in spatial integration and spatial specificity are attributable to the second training phase and not the first one.

**Figure 5:**
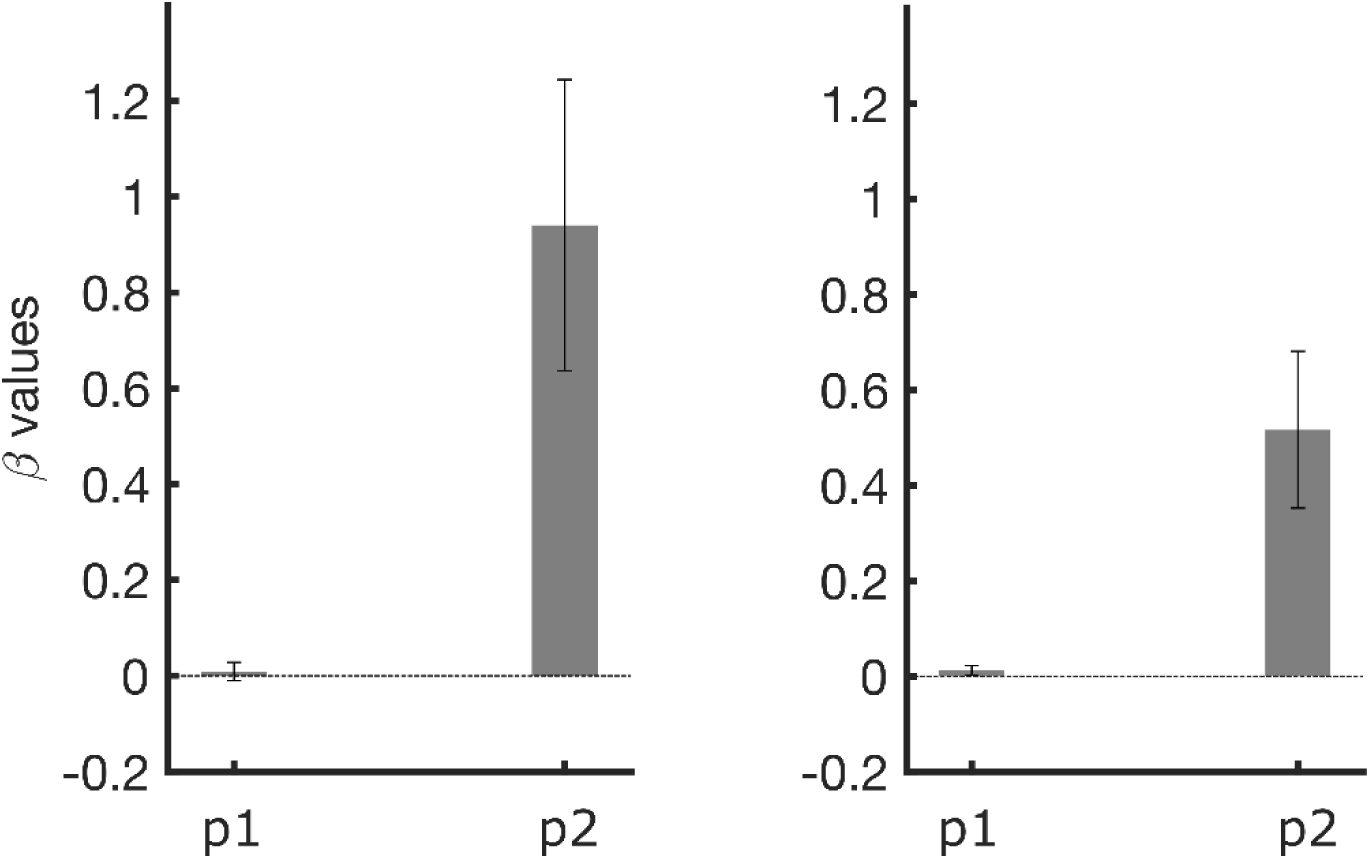
Linear regression analysis. Estimated *β* values for the two training phases (p1 and p2) for both spatial integration (left) and spatial specificity (right). Error bars show SEM.

#### Computational model

Our experiments show that training with a complex stimulus yields a consistent pattern of effects on performance with simpler stimuli: Performance decreases for stimuli at the trained location and increases for stimuli far from the trained location. Likewise, performance decreases for stimuli smaller than the trained stimulus, but increases for stimuli larger than the trained stimulus. We hypothesize that this pattern of changes results from the hierarchical structure of the visual pathways and from the reweighting of neural inputs that occurs during perceptual learning.

To test this idea, we built a computational model comprised of two components. The first approximated the stimulus processing in cortical areas most likely to be involved in our experiments, namely MT and MST. A second component of the model implemented a decoding mechanism that could perform reweighting in response to exposure to different stimuli.

The first component of the model was based on a previously published model of neuronal selectivity in MT and MST that had been statistically validated with real neural recordings (Cui et al., 2013; Mineault et al., 2012). Because neurophysiological evidence suggests that training has little or no effect on the tuning of neurons in these areas (Law & Gold, 2008; Liu & Pack, 2017) we fixed this component and constrained plasticity to occur in the second component, which was meant to function as a downstream decoding mechanisms (Law & Gold, 2008). To this end, we used an adaptive decoder (Jacobs, 2009) that performed an optimal readout of sensory information to perform the motion discrimination tasks used in the current experiments. As in previous work (Dosher et al., 2013), the decoder had access to neurons at multiple stages along the sensory processing hierarchy (Figure 6a).

Figure 6b and 6c show example tunings of simulated MT (Figure 6b) and MST (Figure 6c) neurons for both RDK (left panel in Figure 6b-c) and optic flow (right panel in Figure 6b-c) stimuli at eight positions across the visual field. The tuning curves are plotted as tuning mosaics, in which each mosaic color-codes the firing rate of the neuron evoked by one out of eight possible motions in each motion type (i.e. different directions of RDK motion or different types of optic flow). Orange colors indicate high firing rates, and blue colors indicate low firing rates. The eight tuning mosaics (eight circles in each panel) show the tunings of the neurons for eight different positions in the visual field. For example, the MT neuron shown in Figure 6b has a receptive field centered at the upper left quadrant, and is tuned to downward motion (270° motion direction). This MT neuron, however, is not specifically tuned to optic flow, as it responds maximally to both expanding radial and clockwise circular motions at different positions, consistent with real MT neurons (Lagae et al., 1994). The MST neuron in Figure 6c, on the other hand, has a very large receptive field: it responds with consistent selectivity for spiral optic flow stimuli at most locations (Figure 6c-right), as is true of many MST neurons (Graziano, Andersen, & Snowden, 1994; Lagae et al., 1994). The same MST neuron is also weakly tuned to upward RDK motion (Figure 6c-left).

Following our protocol in Experiment 2, we implemented two training phases in our model: the first used simple RDK motion (leftward/rightward motion discrimination), and the second used optic flow (expansion/contraction radial motion discrimination). In both phases, the training stimulus had a fixed size (diameter = 5°), and was located in the upper left quadrant (*x* = −5°, *y* = −5°). Figure 7a shows the learning curves of the model for the two training phases, demonstrating the improved sensitivity of the model after about 500 iterations. Our hypothesis predicts that training on the simpler stimulus (here RDK) should increase the readout weight of MT neurons, while training on the more complex stimulus (here optic flow) should increase the readout weight of MST neurons. As a measure of MT and MST readout weights, we quantified the ratio of the final pool of sensory neurons that belonged to MT or MST population at the end of each training phase. In Figure 7b the readout weights of MT (in blue) and MST (in green) are shown for the end of each phase. Consistent with our prediction, after training on RDK, the MT readout weight was larger than the MST readout weight (0.9 ± 0.04 *vs*. 0.14 ± 0.05), while this pattern reversed after training on optic flow (0.1 ± 0.04 *vs*. 0.086 ± 0.05).

**Figure 7:**
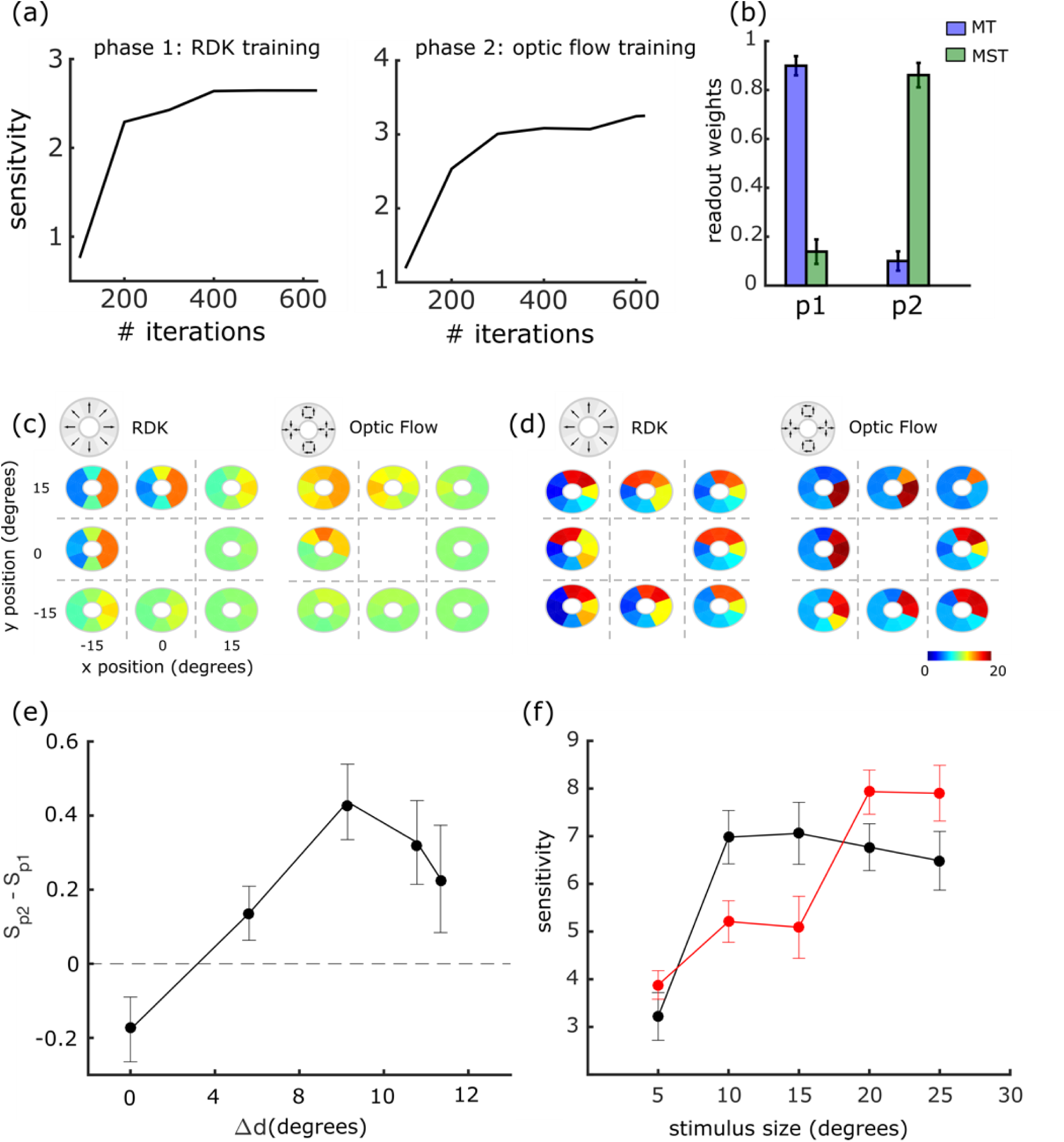
Results of the computational model. (a) Learning curves for the two training phases. (b) The average readout weights of the MT (blue) and MST (green) populations at the end of the first (p1) and second (p2) training phases. (c)-(d) The tuning mosaics of the neurons with the largest readout weight after the first (c) and the second (d) training phases for both RDK (left panel in both (c) and (d)) and optic flow (right panel in both (c) and (d)). (e) The change in the sensitivity of the model as a function of distance from the first to the second training phases. (f) The change in the sensitivity of the model as a function of the stimulus size after RDK (black) and optic flow (red) trainings. Error bars show SEM.

The tunings of the neurons with the largest readout weights at the end of each training phase are shown in Figure 7c-d for both RDK and optic flow stimuli. The MT neuron (Figure 7c) is strongly tuned to rightward motion, an optimal readout for the leftward/rightward RDK training task (Butts & Goldman, 2006). In contrast, the MST neuron (Figure 7d) is strongly tuned to expansion motion, which is the optimal readout for the radial motion training task in the second phase. This result shows that the adaptive decoder was capable of finding the most optimal type of neuron for each training task.

To simulate the specificity tests in our psychophysics experiments, at the end of each training phase, we measured our model’s sensitivity on the simpler stimulus (here RDK) at different distances from the training location. Figure 7e shows the changes in the sensitivity of our model’s output from the first to the second training phases as a function of distance from the trained location. Similar to our psychophysical observations (Figures 2e and 3e), sensitivity increased at locations more distant from the trained location, while it dropped at the trained location. This difference between the trained and the untrained locations can be explained by the differences in the tunings of the MT and MST neurons in Figure 7c-d. The MST neuron (Figure 7d) has a larger receptive field that encompasses more distant locations. Reading out from this neuron at the end of the second training phase improved sensitivity at locations that the receptive field of the MT neuron (Figure 7c) could not reach. However, the suboptimal tuning of the MST neuron for the RDK task compared to the MT neuron (both in terms of receptive field size and direction tuning), explains the observed drop in the performance at the trained location. Thus, in our model the improved sensitivity at a larger spatial scale is necessarily associated with a decrease in the precision of coding local features. This trade-off is reflected in the difference of sensitivity changes between the trained and untrained locations in Figure 7e. Measuring the model’s sensitivity with different stimulus sizes revealed the same trade-off: As shown in Figure 7f, after training with optic flow the sensitivity of the model to large stimuli increased while the sensitivity decreased for small stimuli.

## Discussion

Visual perceptual learning is often specific to the features of the training stimulus, limiting the utility of standard training procedures (Dosher & Lu, 2017). As a result, a variety of different factors have been shown to affect specificity; these include task difficulty (Ahissar & Hochstein, 1997), task complexity (Green, Kattner, Siegel, Kersten, & Schrater, 2015), and the duration of training (Jeter et al., 2009). In this paper, we have shown that training with more complex visual stimuli is associated with lower specificity, even when the task structure is unchanged (Figures 3 and 4). Such decreases in specificity were always accompanied by an increase in psychophysical spatial integration. This trade-off between local specificity and global integration mirrors the gradual increase in the size of the receptive fields along the hierarchy of the visual system. Our findings in this study complement previous studies of visual perceptual learning by suggesting that the stimulus tuning of different visual areas is an important principle that guides the reweighting process.

### Relationship to previous literature

#### Reweighting models of visual perceptual learning

Previous models of visual perceptual learning suggested that the improved performance in perceptual learning is due to optimal reweighting of the sensory neurons for the training task (Dosher & Lu, 1998; Jacobs, 2009; Law & Gold, 2009; Talluri, Hung, Seitz, & Series, 2015). Dosher and colleagues (Dosher et al., 2013; Dosher & Lu, 1998; Petrov, Dosher, & Lu, 2005) showed that a Hebbian learning mechanism that reweights sensory neurons could capture the improvement acquired through perceptual learning with no need for retuning the sensory representations. Importantly, an extended version of their model (Dosher et al., 2013), made use of two stages of visual processing, with larger receptive fields assigned to the higher area: Increased readout weight of the higher sensory stage led to less specificity. This work showed how different training tasks could differentially engage high-level cortical structures, though it did not examine the influence of stimulus complexity.

#### Training stimulus and the specificity of perceptual learning

The relationship between different stimuli and the specificity of visual perceptual learning has been studied previously (Ahissar & Hochstein, 1997; Jeter et al., 2009). Namely, training on more difficult tasks leads to higher specificity (Ahissar & Hochstein, 1997), and these results can be understood as an increase in the contribution of lower visual areas comprised of neurons with narrower tuning and smaller receptive fields. However, it has been argued that *difficulty* cannot be considered as a general feature of a training stimulus, as for different observers and with different stimuli, difficulty could be interpreted differently (Jeter et al., 2009). Therefore, more objective characteristics of visual stimuli need to be studied for understanding the relationship between visual stimuli and the specificity of perceptual learning.

A concept of stimulus complexity akin to the one used in the current study was employed by McGovern et al. (2012) to study the relationship between training stimuli and the specificity of perceptual learning. They evaluated the transfer of learning between different training stimuli with different levels of complexity (McGovern et al., 2012), and observed that transfer was greatest between the stimuli that were closest to each other in terms of complexity. These results are reminiscent of our findings that training with translation RDKs can transfer to DGs (Figure 3) and that training with complex optic flow can transfer to translation RDKs (Figure 4). We did not examine the transfer of learning across stimuli of very different complexity (i.e. between DGs and optic flow), but the results of McGovern et al. (2012) suggest that such transfer would be limited.

One difference between our study and the work of McGovern et al. (2012) is the structure of the training. The latter trained separate groups on different tasks and performed group-wise comparisons, while we had observers complete different training regimens sequentially. In this regard, our approach is similar to that of (Fahle, 1997) who trained observers on a series of tasks and found no transfer in learned improvements. Indeed, observers showed worse performance on one task after training on a second task, which is similar to our finding (Figures 3 and 4) that observers exhibited inferior discrimination for DGs and RDKs at the trained location and size after training on more complex stimuli. The overall picture that emerges is one of a limited scope for learning: Improvements in one domain (space, features) often are accompanied by deterioration of performance in another domain. In some ways, this may not be surprising, as the system has been optimized by evolution and development for performance on real-world visual environments, which contain a broad range of different stimulus features. Nevertheless, our results suggest that in cases where the goal is to improve specific visual capacities, the design of optimal training paradigms can benefit from knowledge of the structure and function of the visual cortex.

Our experimental design in this paper was in some ways similar to the double-training paradigm, which has been shown to decrease specificity in many perceptual learning tasks (Xiao et al., 2008; Zhang et al., 2010). While it seems likely that richer training paradigms, with a greater variety of stimulus types and possible responses, lead to less specificity in learning, this principle does not account for the pattern of results that we have reported. In particular, observers’ performance on measurements of spatial specificity were closely related to the most recently completed training phase. Thus in Experiment 1 we reported a stimulus-dependent difference in specificity after a single training phase (Figure 3f); after Experiment 2, we found that changes in spatial integration could be reversed with further training (Figure 4g); and in both experiments, the outcomes in terms of spatial specificity and integration were predictable from the second phase of training and not the first (Figure 5). In total, while some effects of performing multiple training phases could have been present in our data, the bulk of the observations could be attributed to the stimuli used in individual training phases.

#### Adaptive contribution of sensory neurons in perception

The reweighting model, as well as our results in this study, presume that the contribution of a visual area in a single perceptual task can change adaptively to accommodate different task requirements. This assumption has been supported by previous electrophysiology and neuroimaging studies. In a monkey electrophysiology study, Liu and Pack (2017) showed that training with RDKs increases the contribution of MT to DG motion discrimination. In humans, using transcranial magnetic stimulation (TMS) and fMRI, Chen et al. (2016) showed that the contribution of area V3A in low coherence RDK task can change after RDK training (Chen et al., 2016). Particularly, it was shown that before perceptual learning, V3A and MT were involved in high- and low-coherence motion discrimination, respectively. In contrast, after perceptual learning, V3A became responsible for both conditions (low- and high-coherences). They used TMS inactivation to evaluate the involvement of these visual areas in the tasks pre- and post-training. These results could appear to be at odds with our results, given that their RDK training increased the contribution of a lower visual area (V3A in their case) instead of a higher one. However, a closer look at their training task can resolve this apparent contradiction. In (Chen et al., 2016), during the training, observer performance was measured in terms of the threshold difference between two motion directions that were discriminated. Therefore, throughout the training, the observers were encouraged to distinguish smaller and smaller direction differences. This training criterion does not encourage using larger receptive fields, and actually, V3A with smaller receptive fields than MT could be a more optimal readout option (Jeter et al., 2009). In our study, however, we assessed the observers’ performance with and adaptive coherence threshold, which would require larger spatial integration, and hence, sensory neurons with larger receptive fields.

### Implications for visual rehabilitation in cortical blindness

Visual rehabilitation training for patients with cortical blindness has been an important clinical application of visual perceptual learning. Traditional training protocols have not shown consistent success in recovering the visual abilities of such patients (Pollock et al., 2011). However, recent studies showed that training with RDK stimuli can lead to partial recovery of patients’ visual abilities (Das et al., 2014; Huxlin et al., 2009). Indeed, compared to training with DG, the improvement transfers more to other motion directions and other visual tasks (Das et al., 2014). However, the obtained improvements were still quite limited to the trained location, and retraining was needed to extend the acquired recovery across the scotoma.

Our results suggest that transfer across space can be obtained if we can increase the readout weight of higher visual areas that have the largest receptive fields. Our work suggests that optic flow motion discrimination might increase spatial integration even more than RDKs, presumably by increasing the readout weight of area MST. However, as shown in our results, this improved spatial transfer comes at the cost of sensitivity to small visual stimuli (Figure 4). This trade-off might be satisfactory if the primary goal is to restore functions such as navigation, which benefits from spatial integration.

### Implications for artificial neural networks

The recent success of deep artificial neural networks (DNN) in machine vision tasks, such as object recognition and image classification (Krizhevsky, Sutskever, & Hinton, 2012), has been attributed to the deep hierarchical representation formed across different layers of these models (Bengio, Courville, & Vincent, 2013; LeCun, Bengio, & Hinton, 2015). In the simplest architecture of these models, neurons from each layer project to the immediate next layer, and only the last layer feeds information to the decoder for the decision-making task. The tuning parameters of the neurons across the DNN layers are optimized based on the training task. As shown previously, these deep hierarchical representations also resemble the hierarchical visual representation across different areas of the ventral visual pathway (Yamins et al., 2014).

Because of this strictly hierarchical architecture, only the sensory representation of the deepest layer is accessible to the decoder in many DNNs. This architecture is quite different from that of the primate visual cortex, which exhibits numerous “skip connections”, which bypass hierarchical layers (Felleman & Van Essen, 1991; Tripp, 2019). This is important, since inactivation of the hierarchical flow of information at one stage does not necessarily impair the perceptual output (Liu and Pack, 2017). From a computational standpoint, skip connections could simplify the “credit assignment problem”, wherein errors in the output lead to a difficult search for appropriate synaptic changes in the network. With skip connections, the relationship between low-level representations and the final output can be shallow, simplifying the learning rule. In addition to their recent success in the deep learning applications (He, Zhang, Ren, & Sun, 2016; Huang, Liu, Van Der Maaten, & Weinberger, 2017), architectures with skip connections have also shown to be important in forming latent representations that are most similar to those in the visual system (Schrimpf et al., 2018). Thus, although DNNs have already shown impressive results (Wenliang and Seitz, 2018), an important future direction will be to explore the role of skip connections in visual perceptual learning (Bakhtiari, 2019).

## Acknowledgements

This work was supported by grants from NSERC and the Canada First Research Excellence Fund to C.C.P. We thank Dr. Aaron Seitz for helpful comments on a previous version of the manuscript.

